# Characterization of the impact of daclizumab beta on circulating natural killer cells by mass cytometry

**DOI:** 10.1101/865477

**Authors:** Thanmayi Ranganath, Laura J. Simpson, Christof Seiler, Anne-Maud Ferreira, Elena Vendrame, Nancy Zhao, Jason D. Fontenot, Susan Holmes, Catherine A. Blish

**Author notes:** These authors contributed equally (Thanmayi Ranganath, Laura J. Simpson).

## Abstract

Daclizumab beta is a humanized monoclonal antibody that binds to CD25 and selectively inhibits high-affinity IL-2 receptor signaling. As a former treatment for relapsing forms of multiple sclerosis (RMS), daclizumab beta induces robust expansion of the CD56^bright^ subpopulation of NK cells that is correlated with the drug’s therapeutic effects. As NK cells represent a heterogeneous population of lymphocytes with a range of phenotypes and functions, the goal of this study was to better understand how daclizumab beta altered the NK cell repertoire to provide further insight into the possible mechanism(s) of action in RMS. We used mass cytometry to evaluate expression patterns of NK cell markers and provide a comprehensive assessment of the NK cell repertoire in individuals with RMS treated with daclizumab beta or placebo over the course of one year. Treatment with daclizumab beta significantly altered the NK cell repertoire compared to placebo treatment. As previously reported, daclizumab beta significantly increased expression of CD56 on total NK cells. Within the CD56^bright^ NK cells, treatment was associated with multiple phenotypic changes, including increased expression of NKG2A and NKp44, and diminished expression of CD244, CD57, and NKp46. While the changes were less dramatic, CD56^dim^ NK cells responded distinctly to daclizumab beta treatment, with higher expression of CD2 and NKG2A, and lower expression of FAS-L, HLA-DR, NTB-A, NKp30, and Perforin. Together, these data indicate that the expanded NK cells share features of both immature and mature NK cells. These findings show that daclizumab beta treatment is associated with unique changes in NK cells that may enhance their ability to kill autoreactive T cells or to exert immunomodulatory functions.

## Introduction

Natural killer (NK) cells are innate lymphocytes that are best known for the eponymous function: that of killing other cells. Yet NK cells also play a critical role in immune regulation by secreting cytokines that influence the character of the immune response. NK cell function is controlled by signals received through activating, inhibitory and cytokine receptors (Vivier et al. 2011). Activating receptors, examples of which include the natural cytotoxicity receptors NKp30, NKp44, and NKp46, the C-type lectin receptors NKG2D and NKG2C, and certain classes of activating Killer-cell Immunoglobulin-like Receptors (KIRs), generally sense stress on the target cell and promote NK cell activation. Inhibitory receptors, including most KIRs, NKG2A, and LILRB1 (CD85j), generally recognize Major Histocompatibility Class (MHC) I receptors and dampen NK cell responses to normal healthy cells. Broadly, NK cells are divided into two major classes. In the blood, the mature CD56^dim^ NK cells are the predominant subset and have potent cytotoxic activity, while the relatively immature CD56^bright^ NK cells are generally present at <10% and primarily secrete cytokines. Recent studies have demonstrated significant heterogeneity in the human NK cell repertoire, with a wide range of NK cell subsets expressing different combinations of these activating and inhibitory receptors (Horowitz et al. 2013; Strauss-Albee and Blish 2016; Strauss-Albee et al. 2015; Wilk and Blish 2018; Vosshenrich and Di Santo 2007).

NK cells also express a wide range of cytokine receptors making them extremely responsive to cytokine stimulation. NK cells undergo dramatic shifts in phenotype and function in the presence of cytokines such as IL-2, IL-12, IL-15, and IL-18, singly and in combination (Vendrame et al. 2017; Romee et al. 2012; Fehniger and Cooper 2016). IL-2 plays a particularly critical role in activating NK cells by binding to the low affinity IL-2 receptor, a heterodimer of CD122 (IL-2Rβ) and CD132 (IL-2R□, otherwise known as the common gamma chain). In general, NK cells do not express the high affinity IL-2 receptor, CD25 (IL-2R□). The CD56^bright^ NK cell subset expresses much higher levels of CD122 than the CD56^dim^ subset (Bielekova et al. 2006; Martin et al. 2010; Caligiuri et al. 1990).

Daclizumab is a humanized monoclonal antibody that irreversibly blocks CD25, preventing signaling through the high affinity IL-2R while increasing IL-2 bioavailability to bind to the low affinity receptor (reviewed in (Cohan et al. 2019; Bielekova 2019)). Due to the complex roles of IL-2 *in vivo*, daclizumab induces several immunological changes, including inhibition of T cell activation, reduction in the frequency and survival of regulatory T cells, and expansion of CD56^bright^ NK cells (Cohan et al. 2019; Bielekova 2019). It was originally developed as an intravenous treatment for several disease indications, including the prevention of transplant rejection and the treatment of severe uveitis and T cell leukemia (Cohan et al. 2019; Bielekova 2019). Later a subcutaneous form (daclizumab beta) was developed and approved for the treatment of relapsing forms of multiple sclerosis (RMS) due to its beneficial effects including reduction in lesion size and slowed disease progression (Bielekova et al. 2006, 2009; Elkins et al. 2015; Wynn et al. 2010; Elkins et al. 2012); (Gold et al. 2013). In these initial trials, daclizumab beta treatment was associated with adverse events including cutaneous reactions, malignancies, infections, and transaminase elevations, though these were not sufficiently severe to preclude approval. In 2018, daclizumab beta was voluntarily withdrawn from the market due to the nature and complexity of adverse events associated with the drug and limited number of patients treated, which presented challenges in further characterizing its evolving benefit/risk profile. Subsequently, cases of immune-mediated encephalitis were confirmed as adverse drug reactions that can be related to treatment with daclizumab beta.

While no longer used therapeutically, a better understanding of the effects of daclizumab beta may provide insight into the pleiotropic effects of IL-2 in the setting of RMS. Surprisingly, the beneficial effects of daclizumab beta treatment were linked not to changes in T cell function, but instead were strongly correlated with expansion of CD56^bright^ NK cells (Bielekova et al. 2006; Elkins et al. 2015; Cohan et al. 2019; Bielekova 2019). Although CD56^bright^ NK cells generally have poor cytotoxic activity, the daclizumab beta-expanded CD56^bright^ NK cells could kill activated, autologous CD4^+^ T cells, potentially driving the therapeutic effect by eliminating autoreactive T cells (Bielekova et al. 2006; Jiang et al. 2011). This study was undertaken to provide a better understanding of the effects of daclizumab beta on circulating NK cells *in vivo*.

## Materials and Methods

### Study subjects

Cryopreserved peripheral blood mononuclear cells (PBMCs) from daclizumab beta-treated and placebo-treated individuals living with RMS were chosen from the Biogen SELECT (NCT00390221) and DECIDE (NCT01064401) studies (Kappos et al. 2015; Gold et al. 2013). Subjects were treated subcutaneously with 150mg daclizumab beta every 4 weeks for 52 weeks. For the placebo and the treatment cohort, we received de-identified PBMCs at 3 timepoints: Baseline, Week 24 and Week 52. For the healthy donor cohort, leukoreduction system chambers from anonymous donors were purchased from the Stanford Blood Bank. PBMCs were isolated by Ficoll density gradient centrifugation and then cryopreserved in fetal bovine serum (FBS) with 10% dimethyl sulfoxide (DMSO). We had 17 healthy donors, 17 placebo and 30 daclizumab beta treated individuals. As part of their initial enrollment, all subjects provided written informed consent. The studies were approved by the relevant central and local ethics committees and were conducted in accordance with the International Conference on Harmonisation guidelines for Good Clinical Practice and the principles of the Declaration of Helsinki.

### Antibody Conjugation, Mass Cytometry Staining and Data Acquisition

Antibodies for mass cytometry were conjugated to heavy metals using MaxPar® ×8 labeling kits (Fluidigm) as described (Kay, Strauss-Albee, and Blish 2016). To ensure antibody stability over time, the antibody panel was lyophilized into single-use pellets prior to use (Biolyph). PBMCs were thawed at 37°C in RPMI-1640 media (supplemented with 10% FBS, L-glutamine, and Penicillin-Streoptomycin-Amphotericin) with benzonase. NK cells were purified by magnetic bead isolation via negative selection (Miltenyi, cat. 130-092-657) and stained with the NK cell antibody panel (Table S1) as previously described (Vendrame et al. 2019; McKechnie et al. 2019; Strauss-Albee et al. 2015). Cells were resuspended in 1x EQ Beads (Fluidigm) for normalization before acquisition on a Helios mass cytometer (Fluidigm).

### Data analysis

The open source statistical software R (https://www.r-project.org/) was used for all statistical analyses (R Core Team 2014). Signal intensities were transformed using the hyperbolic sine transformation (asinh function) prior to statistical analysis, with cofactor equal to 5, to account for heteroskedasticity. We used the custom-made package *CytoGLMM* (Seiler et al. 2019; Kronstad et al. 2018) to identify markers predictive of a given sample type while taking into account the subject effect. To this end, this package uses a generalized linear mixed model with paired comparison (used for analyses of the same individual over time) and generalized linear model with bootstrap resampling (for cross-sectional comparisons between daclizumab beta- and placebo-treated individuals). Using the empirical marker distribution, the model generates the log-odds that the expression of a given marker is predictive of the sample type (for example, drug-treated vs. placebo-treated) with the 95% confidence intervals. For paired comparisons, we computed *p*-values using the asymptotic theory implemented in R package *mbest* (Perry 2017). For unpaired comparisons, we computed *p*-values by inverting the percentile bootstrap confidence intervals and assuming two-sided intervals with equal tails (Efron and Tibshirani 1993). To correct for multiple comparisons, the Benjamini-Hochberg method controlling the False Discovery Rate at level 0.05 was used, which is conservative as it assumes independence of markers. For paired analysis on daclizumab beta-treated individuals over time, where the response variable was timepoint, we used all cells from each donor. For unpaired analysis using the bootstrap, where the response variable was daclizumab beta or placebo treatment, we used 1,000 cells from each sample for the total NK cell and CD56^dim^ analyses, and used all cells from each sample for CD56^bright^. There were fewer than 1,000 CD56^bright^ NK cells in most samples except for daclizumab beta treated individuals. The number of subjects used for each analysis is specified in the figure legends.

### UMAP visualizations

The Uniform Manifold Approximation and Projection (UMAP) algorithm was used as a visualization and dimensionality reduction technique for our CyTOF data (McInnes, Healy, and Melville 2018; Becht et al. 2018). The *uwot* R package provides an implementation of UMAP and was used with a minimum distance set to 0.1 and nearest neighbors set to 3. The UMAP loadings were visualized using Cytobank. Separate analyses were performed on total NK cells and CD56^bright^ NK cells, including both placebo and drug treatment at three different timepoints. All markers in Table S1 were used excluding markers used for gating (CD3, CD19, CD33, CD14, CD56, CD4), and markers with extremely low or nonspecific staining (FcR□, Ki-67, KIR2DS2, CXCR6, PD1).

### Data availability

The dataset generated and analyzed for this study can be found in FlowRepository (accession number XXX).

## Results

### Characteristics of study population

For this study, individuals living with RMS received 150mg daclizumab beta or placebo subcutaneously every 4 weeks for 52 weeks. The demographics of the healthy controls, placebo-treated, and daclizumab beta-treated groups are given in Table 1. Peripheral blood samples for our research study were taken from both the SELECT and DECIDE trials (Kappos et al. 2015; Gold et al. 2013).

**Table 1:** Demographics of the study population.

| Group | Total | Pct. Female | Age, years: median (range) |
| --- | --- | --- | --- |

|  | N |  |  |
| --- | --- | --- | --- |
| **Healthy** | **12** | **50%** | **52.5 (27-82)** |
| **Placebo** | **17** | **70.5%** | **34 (21-45)** |
| **Daclizumab beta** | **30** | **90%** | **32 (19-52)** |

### Characterization of the NK repertoire

Frozen peripheral blood samples from baseline (pre-treatment), 24 weeks post-treatment initiation, and 52 weeks post-treatment initiation were obtained for this study. Purified NK cells from each sample were stained for mass cytometry using a panel of 41 antibodies conjugated to heavy metals (Table 2, Figure S1A). Total NK cells, CD56^bright^ and CD56^dim^ NK cells were analyzed by gating in FlowJo (Figure S1B). The UMAP algorithm was used to visualize NK cells from each group at each timepoint. The spatial organization of well-known NK cell subsets in placebo recipients at baseline are demonstrated in Figure S1C, and are preserved between groups.

### Daclizumab beta induces higher frequency of CD56^bright^ NK cells

The *CytoGLMM* R package was used to identify which NK markers predicted daclizumab beta treatment compared to placebo. This generalized linear model with bootstrap resampling allows for identification of markers that predict a given outcome, while controlling for inter-individual variability. The model takes into account the full distribution of the marker measurements (rather than a single summary measure such as mean signal intensity) and yields the log-odds with which that marker predicts the outcome, with 95% confidence intervals. Among total NK cells at 24 weeks, NKp30, NTB-A, and CD2 expression predicted daclizumab beta treatment, while NKG2D, CD244, TIGIT, FAS-L, and KIR2DL5 predicted placebo treatment (Figure 1A). After controlling for multiple comparisons, these changes were not statistically significant. At 52 weeks, among the total NK cell population, CD56, NKp30, TACTILE, NKp44, and NTB-A predicted daclizumab beta treatment, while CD244, CD69, TIGIT, NKp46, and CD57 predicted placebo treatment (Figure 1A), but these changes were not significant after correction for multiple comparisons.

**Figure 1.**
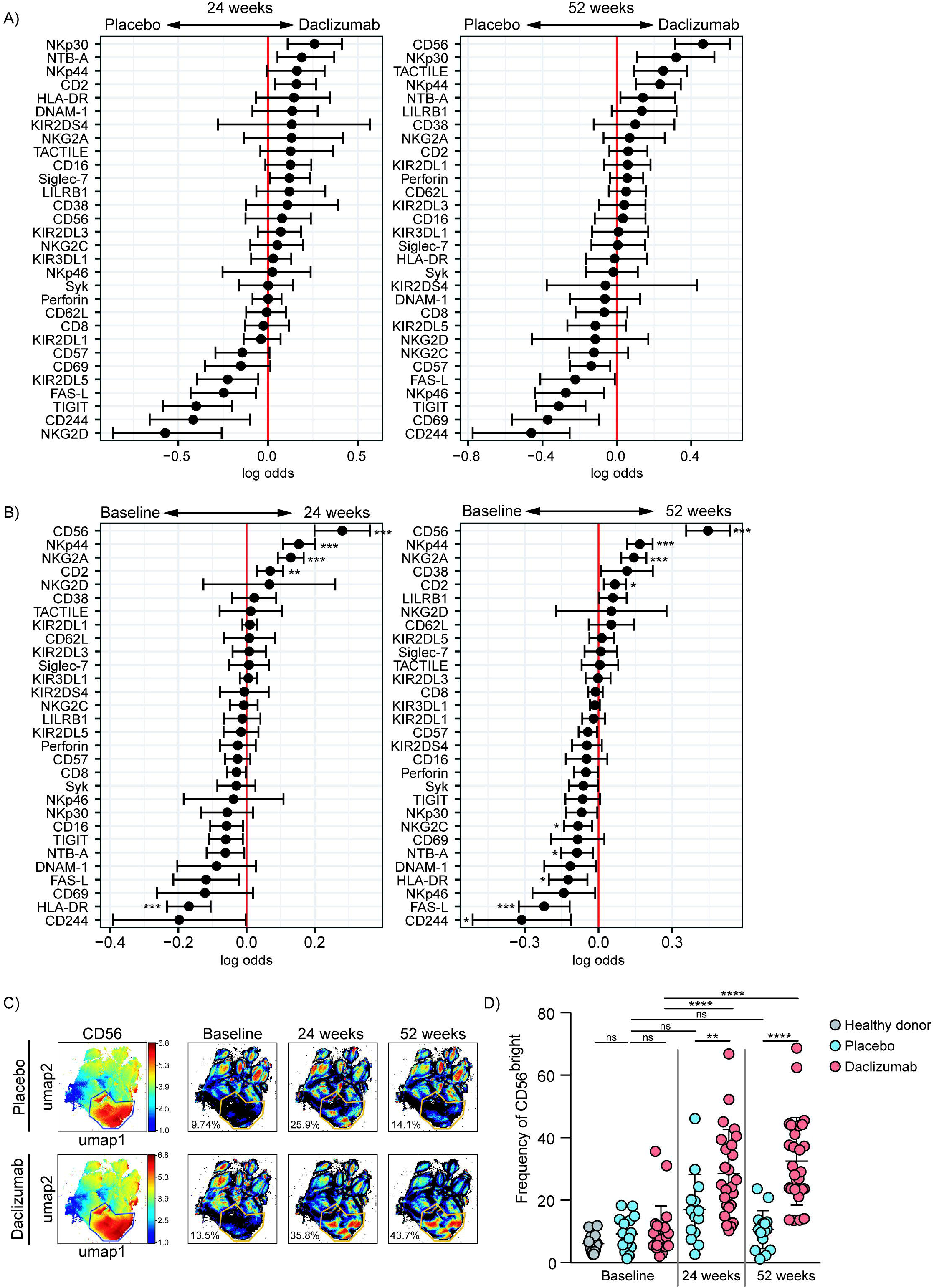
Daclizumab beta induces CD56^bright^ NK cells with a distinct phenotype. **(A)** A generalized linear model with bootstrap resampling was used to identify NK markers predictive of daclizumab beta- and placebo-treated individuals at 24 (left) and 52 (right) weeks of treatment. Log-odds are logarithm of ratios of the probability that a cell belongs to each treatment group. An increase in the parameter coefficient corresponds to the strength of the classification power, with the 95% confidence interval represented by the line line surrounding the point estimate. Total NK cells were used, with subsampling to 1000 cells per individual. Placebo: 24 weeks, n = 14; 52 weeks, n = 16. Daclizumab beta: 24 weeks, n = 25; 52 weeks, n = 27. **(B)** A generalized linear mixed model with paired comparison was used for analyses of the same individual over time, comparing baseline and 24 weeks of daclizumab beta treatment (left), and baseline and 52 weeks of daclizumab beta treatment (right). Total NK cells were used with no subsampling. Daclizumab beta baseline vs. 24 weeks, n = 21. Daclizumab beta baseline vs. 52 weeks, n = 22. **(C)** UMAP visualization of all NK cells from placebo and daclizumab beta treatment groups. Panels on the left are colored by CD56 expression, with a gate drawn around the CD56^bright^ population. Baseline, 24 weeks, and 52 weeks samples are shown colored by density, with the same CD56^bright^ gate. Percentages represent the number of cells that fall within the CD56^bright^ gate. **(D)** The frequency of CD56^bright^ NK cells for each individual are shown as a percentage of total NK cells. Healthy donors (n = 16, grey), placebo treatment (baseline, n = 16; 24 weeks, n = 14; 52 weeks, n = 16; blue), daclizumab beta treatment (baseline, n = 22; 24 weeks, n = 25, 52 weeks, n = 27; pink). *Adjusted p-value < 0.05, **adjusted p-value < 0.01, ***adjusted p-value < 0.001, ****adjusted p-value < 0.0001. Adjusted p-values calculated on generalized linear mixed model in (B) using Benjamini-Hochberg method with FDR = 0.05. Adjusted p-values in (D) calculated using one-way ANOVA with Sidak’s multiple comparisons test.

Within the daclizumab beta-treated group, most protein expression changes that occurred in total NK cells by 24 weeks were preserved at 52 weeks. CD56, NKp44, NKG2A, and CD2 predicted 24 weeks of treatment, while HLA-DR and FAS-L predicted baseline samples (Figure 1B). With the increased power from these paired comparisons, the changes in CD56, NKp44, NKG2A, CD2, and HLA-DR were significant after correction for multiple comparisons. When comparing baseline and 52 weeks, CD56, NKp44, NKG2A, and CD2 significantly predicted 52 weeks of treatment, while CD244, FAS-L, HLA-DR, NTB-A, and NKG2C significantly predicted baseline samples (Figure 1B).

CD56 was the most significant predictor of daclizumab beta treatment, and predicted both 24- and 52-week samples compared to baseline. Using UMAP visualization, we found an increase in frequency of CD56^bright^ NK cells by 24 weeks of daclizumab beta treatment, which continued to increase at 52 weeks (Figure 1C). This increase in CD56^bright^ NK cells was confirmed by gating in FlowJo (Figure 1D, Figure S1B), which showed that the frequency of CD56^bright^ NK cells was increased in daclizumab beta-treated subjects compared to placebo-treated, and increased from baseline to 24 weeks and 52 weeks. There was no significant difference in frequency of CD56^bright^ NK cells in placebo-treated individuals over time, or in individuals with MS before treatment compared to healthy controls.

### Daclizumab beta alters expression of NK receptors on the CD56^bright^ population compared to placebo

As the CD56^bright^ and CD56^dim^ NK cell subsets are distinct, we next focused solely on the CD56^bright^ NK cells (Figure S1B for gating strategy). The *CytoGLMM* package was used to identify which markers predicted daclizumab beta treatment compared to placebo within the CD56^bright^ population. CD2, NKp30, and Siglec-7 predicted daclizumab beta treatment compared to placebo after 24 weeks of treatment in unadjusted comparison, but were not significant following correction for multiple comparisons (Figure 2A, panel 1). NKp30, Perforin, NKp44, TACTILE, Siglec-7, KIR2DL3, and CD16 predicted daclizumab beta treatment compared to placebo after 52 weeks of treatment in the unadjusted comparison, but was not significant following adjustment for multiple comparisons (Figure 2A, panel 2).

**Figure 2.**
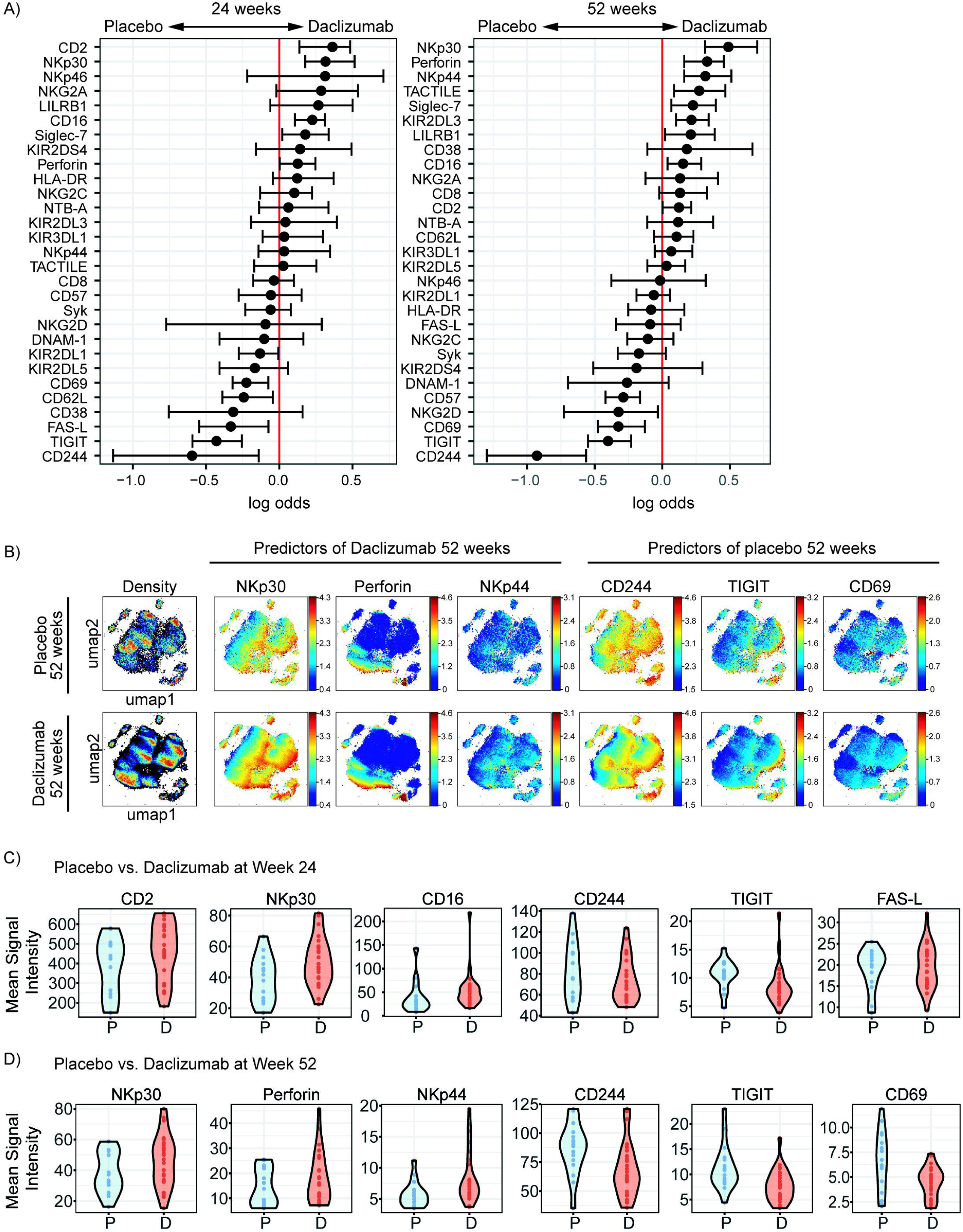
Daclizumab beta alters the CD56^bright^ population in comparison with placebo. **(A)** A generalized linear model with bootstrap resampling was used to identify NK markers on CD56^bright^ NK cells predictive of daclizumab beta- and placebo-treated individuals at 24 (left) and 52 (right) weeks of treatment. CD56^bright^ NK cells were used with no subsampling. Placebo: 24 weeks, n = 14; 52 weeks, n = 16. Daclizumab beta: 24 weeks, n = 25; 52 weeks, n = 27. **(B)** UMAP visualizations of CD56^bright^ NK cells from the placebo (top row) or the daclizumab beta-treated (bottom row) groups at 52 weeks. Leftmost panels are colored by cell density. NKp30, Perforin, and NKp44 were predictors of daclizumab beta treatment. CD244, TIGIT, and CD69 were predictors of placebo treatment. Each plot is colored by marker expression, with color scale consistent between groups but specific for each marker. **(C)** Violin plots of mean signal intensity of CD2, NKp30, and CD16 (predictors of daclizumab beta at week 24), and CD244, TIGIT, and FAS-L (predictors of placebo at week 24). Data points represent the mean signal intensity per individual. Placebo week 24, n = 14. Daclizumab beta week 24, n = 25. **(D)** Violin plots of mean signal intensity of NKp30, Perforin, NKp44 (predictors of daclizumab beta at week 52), and CD244, TIGIT, and CD69 (predictors of placebo at week 52). Data points represent the mean signal intensity per individual. Placebo week 52, n = 16. Daclizumab beta week 52, n = 27.

Expression of each of the top three predictors of 52 weeks of daclizumab beta treatment and the top three predictors of placebo treatment was visualized using UMAP (Figure 2B). In the UMAP projections, the plots show an overall increase in density of CD56^bright^ NK cells at 52 weeks of daclizumab beta treatment (Figure 2B). NKp30 has higher expression in the CD56^bright^ population of the daclizumab beta treated group, and particularly higher expression in the areas of high cell density. Conversely, CD244 predicted placebo treatment, and had higher expression across the CD56^bright^ population of placebo treated individuals.

In order to look at expression of each of the strongest predictors of daclizumab beta or placebo treatment in each individual, the mean signal intensity was calculated at 24 weeks (Figure 2C) or 52 weeks (Figure 2D). There was a modest increase in CD2, NKp30, and CD16 in the daclizumab beta treated individuals compared to placebo at 24 weeks of treatment, and a modest decrease in CD244, TIGIT, and FAS-L. The increase in NKp30 expression became stronger after 52 weeks of treatment (Figure 2D). At 52 weeks, Perforin and NKp44 were also strong predictors of daclizumab beta treatment, and their expression levels were increased in daclizumab beta treated individuals. The decrease in expression of CD244 and TIGIT observed at 24 weeks of daclizumab beta treatment was more dramatic after 52 weeks of treatment. Interestingly, CD69, a marker of early activation, was significantly decreased after 52 weeks of daclizumab beta treatment, suggesting that the CD56^bright^ population is not necessarily more activated.

### Daclizumab beta alters NK receptor expression in the CD56^bright^ population over 52 weeks of treatment

In order to determine what changes in NK receptor expression occurred in response to daclizumab beta in treated individuals over time, the predictors of baseline or 24 weeks of treatment were determined using *CytoGLMM* (Figure 3A). Many NK receptors significantly predicted 24 weeks of daclizumab beta treatment: NKG2A, NKp44, KIR2DL3, CD8, CD38, Siglec-7, FAS-L, Perforin, and CD16 (Figure 3A, panel 1). Interestingly, the top six predictors of 24 weeks of daclizumab beta treatment also predicted 52 weeks of treatment, suggesting that most of the daclizumab beta-induced changes in NK receptor expression occur within the first 24 weeks of treatment and are maintained throughout treatment (Figure 3A, panel 2). CD244, CD57, CD69, NKp46, Syk, DNAM-1, and KIR2DL5 significantly predicted baseline samples compared to both 24 weeks and 52 weeks. TIGIT and NKG2C additionally predicted baseline samples compared to 52 weeks of daclizumab beta treatment.

**Figure 3.**
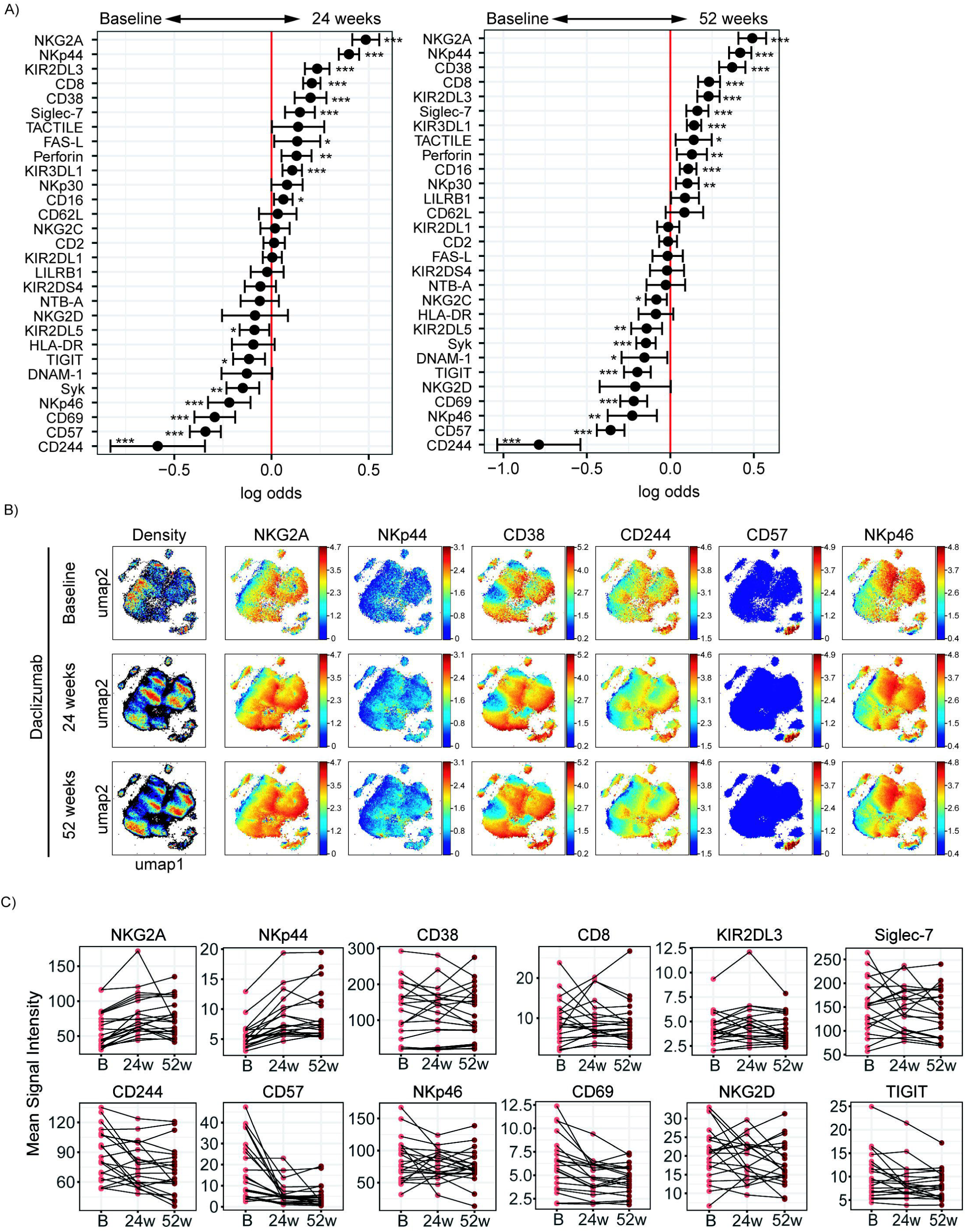
Daclizumab beta-induced NK receptor expression changes on CD56^bright^ NK cells occur in the first 24 weeks. **(A)** A generalized linear mixed model with paired comparison was used for analyses of the same individual over time, comparing baseline and 24 weeks of daclizumab beta treatment (left), and baseline and 52 weeks of daclizumab beta treatment (right). CD56^bright^ NK cells were used with no subsampling. Daclizumab beta baseline vs. 24 weeks, n = 21. Daclizumab beta baseline vs. 52 weeks, n = 22. **(B)** UMAP visualizations of CD56^bright^ NK cells from the daclizumab beta-treated group at baseline (top row), 24 weeks (middle row), and 52 weeks (bottom row). Leftmost panels are colored by cell density. NKG2A, NKp44, and CD38 were predictors of 52 weeks of treatment. CD244, CD57, and NKp46 were predictors of baseline samples. Each plot is colored by marker expression, with color scale consistent between groups but specific for each marker. **(C)** Mean signal intensity of NKG2A, NKp44, CD38, CD8, KIR2DL3, and Siglec-7 (predictors of daclizumab beta at week 52). Mean signal intensity of CD244, CD57, NKp46, CD69, NKG2D, and TIGIT (predictors of baseline samples). Data points represent the mean signal intensity per individual, with lines connecting each individual over time. Only subjects with samples at all three timepoints are shown; n = 20. *Adjusted p-value < 0.05, **adjusted p-value < 0.01, ***adjusted p-value < 0.001. Adjusted p-values calculated using Benjamini-Hochberg method with FDR = 0.05.

UMAP visualizations of each of the top three predictors of 52 weeks of treatment and baseline are shown in Figure 3B. NKG2A significantly increased expression by 24 weeks, and maintained high levels of expression at 52 weeks of treatment, especially in areas of high cell density in the UMAP projections. Conversely, CD244 expression broadly decreased across the CD56^bright^ population by 24 weeks, and remained reduced at 52 weeks. Not all changes in NK receptor expression occurred across the whole CD56^bright^ population. For example, CD57 expression appears unchanged in the UMAP projection colored by CD57; however, when compared with the cell density plot, it becomes clear that the small CD57^+^ island is reduced in cell density by 24 and 52 weeks of daclizumab beta treatment.

The mean signal intensity was calculated for each of the top six predictors of 52 weeks of daclizumab beta treatment and the top six predictors of baseline samples in each individual (Figure 3C). NKG2A and NKp44 expression increased in almost every individual by 24 weeks of daclizumab beta treatment, and tended to stay elevated at 52 weeks. CD38, CD8, KIR2DL3, and Siglec-7 expression increased in some but not all subjects, suggesting that the daclizumab beta-induced CD56^bright^ population is not equivalent in all subjects receiving daclizumab beta. CD244, CD57, NKp46, CD69, and TIGIT expression were significantly decreased by 24 weeks of daclizumab beta treatment in nearly all subjects, while NKG2D expression varied between individuals. These graphs highlight the fact that most of the changes in NK receptor expression observed with daclizumab beta treatment occur early in treatment, and are maintained throughout the course of treatment.

### Daclizumab beta also alters NK receptor expression in the CD56^dim^ population

While the focus of this study was to determine NK receptor expression in the daclizumab beta-induced CD56^bright^ population, it was interesting to find that there were some distinct changes in NK receptor expression observed in the CD56^dim^ population as well. Using *CytoGLMM*, we identified several weak (low log-odds) predictors of daclizumab beta treatment compared to placebo in the CD56^dim^ population in the unadjusted analysis, including NTB-A and CD2 at 24 weeks, and TACTILE, NTB-A, and CD2 at 52 weeks; these findings were not significant after correcting for multiple comparisons (Figure 4A). There were stronger (higher log-odds) predictors of placebo treatment at both 24 and 52 weeks, including NKG2D, TIGIT, FAS-L, KIR2DL5, CD69, CD244, and NKp46 in the unadjusted analysis (Figure 4A). This suggests that daclizumab beta more strongly decreased expression of several NK receptors in the CD56^dim^ population rather than increased.

**Figure 4.**
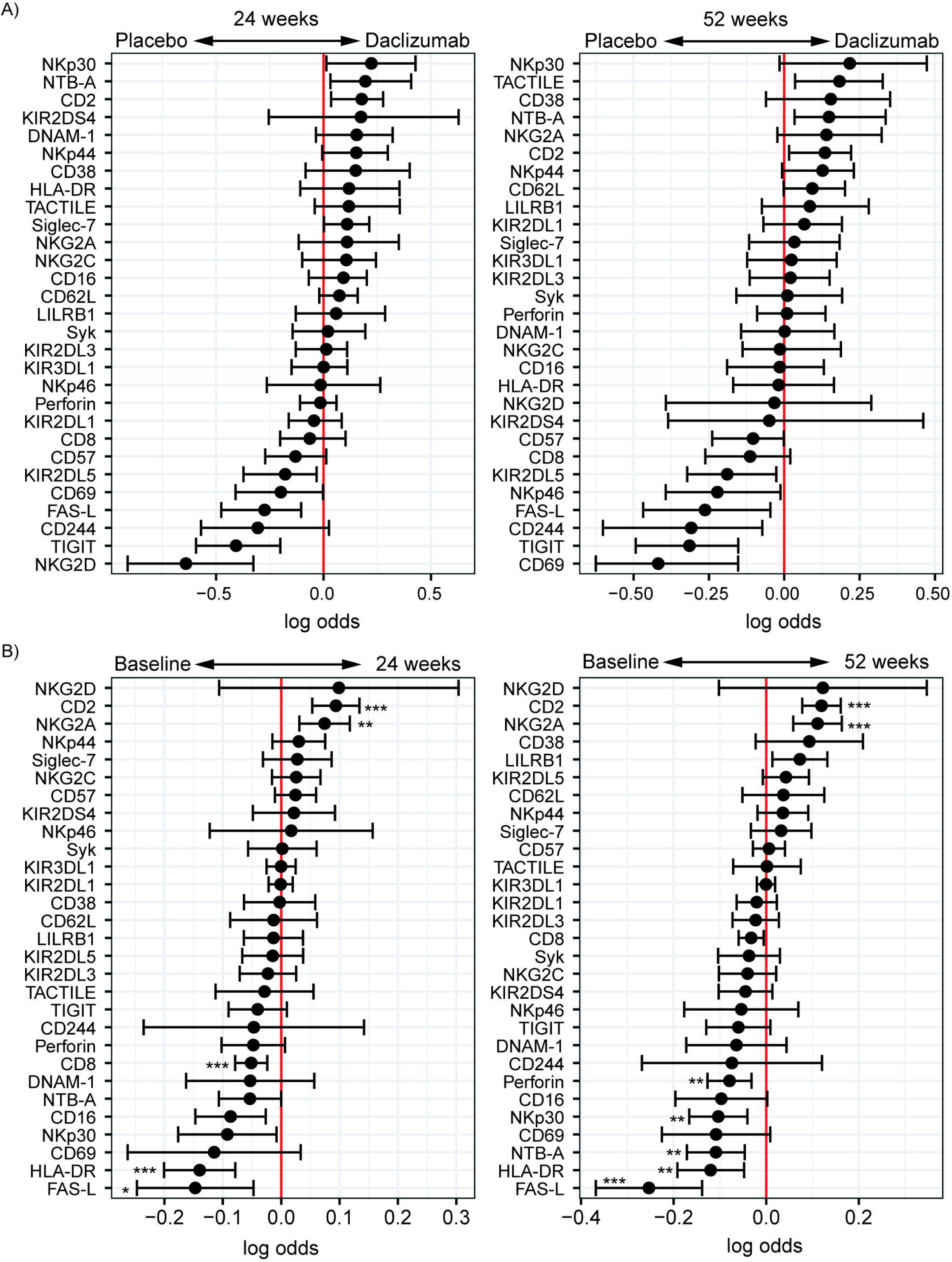
Daclizumab beta changes NK receptor expression on CD56^dim^ cells distinctly from CD56^bright^ cells. **(A)** A generalized linear model with bootstrap resampling was used to identify NK markers on CD56^dim^ NK cells predictive of daclizumab beta- and placebo-treated individuals at 24 (left) and 52 (right) weeks of treatment. CD56^dim^ NK cells were used with subsampling to 1000 cells per individual. Placebo: 24 weeks, n = 14; 52 weeks, n = 16. Daclizumab beta: 24 weeks, n = 25; 52 weeks, n = 27. **(B)** A generalized linear mixed model with paired comparison was used for analyses of the same individual over time, comparing baseline and 24 weeks of daclizumab beta treatment (left), and baseline and 52 weeks of daclizumab beta treatment (right). CD56^dim^ NK cells were used with no subsampling. Daclizumab beta baseline vs. 24 weeks, n = 21. Daclizumab beta baseline vs. 52 weeks, n = 22. *Adjusted p-value < 0.05, **adjusted p-value < 0.01, ***adjusted p-value < 0.001. Adjusted p-values calculated using Benjamini-Hochberg method with FDR = 0.05.

When comparing baseline and 24 or 52 weeks of treatment in daclizumab beta-treated individuals, we identified two significant predictors of treatment: CD2 and NKG2A (Figure 4B). Conversely, FAS-L, HLA-DR, CD8, NTB-A, NKp30, and Perforin predicted baseline samples compared to 24 or 52 weeks. Interestingly, several of the markers that predicted baseline samples in the CD56^dim^ population actually predicted daclizumab beta treatment in the CD56^bright^ population, suggesting that distinct responses to daclizumab beta treatment occur in the CD56^bright^ and CD56^dim^ NK cells.

## Discussion

While daclizumab beta has been removed from the market, understanding of the effects of daclizumab beta may provide insight into the role of NK cells in the pathogenesis of RMS and in the potential of other NK cell-based therapeutic strategies for RMS. Here, we used mass cytometry to profile the NK cells expanded in the setting of daclizumab beta treatment of RMS. As observed previously (Bielekova et al. 2006), daclizumab beta led to a dramatic shift in the NK cell repertoire with an increase in the frequency of CD56^bright^ NK cells and enhanced expression of NKG2A and CD2. Here, we extended the characterization of the phenotype, demonstrating that the expanded CD56^bright^ NK cells had a unique phenotype that was present by 24 weeks of treatment.

Daclizumab beta treatment was associated with enhanced expression of the activation markers CD38 and Perforin, the activating receptors NKp44, TACTILE (CD96), and CD16, and the inhibitory receptors NKG2A, KIR2DL3, and Siglec-7 within the CD56^bright^ population. The enhanced expression of a number of markers of cellular activation and receptors that mediate NK cell activation suggests that the CD56^bright^ NK cells that emerge upon daclizumab beta treatment are primed for responsiveness. At the same time, it is difficult to fully predict the effects of these alterations in receptor expression profiles. While CD56^bright^ NK cells normally express NKG2A, the inhibitory receptor that binds HLA-E, this expression is even higher following daclizumab beta treatment (Figure 3). While enhanced expression of this inhibitory receptor could diminish NK cell responsiveness, it is also possible that these NK cells are ‘educated’ through NKG2A, leading to their enhanced ability to detect ‘altered self’ (Ramsuran et al. 2018; Horowitz et al. 2016). Along similar lines, the increased expression of KIR2DL3 on the CD56^bright^ NK cells in daclizumab beta-treated subjects indicates that at least a subset of these CD56^bright^ NK cells have a more mature phenotype, and could be educated through this KIR to enhance their ability to respond to missing self. Thus, daclizumab beta treatment drives several phenotypic changes in NK cells that could contribute to enhanced NK cell responsiveness.

Daclizumab beta treatment had less effect on the mature CD56^dim^ subset that is generally associated with higher cytolytic activity, yet phenotypic changes still occurred. Similar to CD56^bright^ NK cells, expression of NKG2A and CD2 on CD56^dim^ cells was predictive of daclizumab beta treatment. Interestingly, several markers associated with cellular cytotoxicity, including FAS-L and Perforin, predicted baseline samples among CD56^dim^ NK cells, suggesting that the CD56^dim^ NK cells from daclizumab beta treated subjects may be less efficient at killing. This could partially explain the killing of autoreactive T cells by CD56^bright^ NK in the setting of daclizumab beta treatment (Bielekova et al. 2006).

In general, CD56^bright^ NK cells are thought of as the immature subset of NK cells that do not express KIRs, have limited cytolytic activity, and secrete cytokines. Their expansion following daclizumab beta treatment could therefore improve outcomes in RMS through either immunomodulation by cytokine secretion or by killing autoreactive T cells. Surprisingly, prior work demonstrated that the daclizumab beta-expanded CD56^bright^ NK cells can kill activated autologous CD4^+^ T cells in a granzyme K dependent manner *in vitro* (Bielekova et al. 2006; Jiang et al. 2011), which may well explain the therapeutic effect of daclizumab beta. The fact that there was a strong correlation between CD56^bright^ NK cell expansion and T cell contraction following daclizumab beta treatment supports the idea that the CD56^bright^ NK cells could be limiting disease by eliminating T cells *in vivo* (Bibiana Bielekova et al. 2006). One potential explanation for this surprising cytotoxicity mediated by CD56^bright^ NK cells is that they are not ‘conventional’ CD56^bright^ NK cells, but instead fully mature CD56^dim^ NK cells that upregulated CD56 expression to the point where they were re-classified as CD56^bright^ NK cells. In fact, prior work demonstrates that IL-2 increases CD56 expression on NK cells (Vendrame et al. 2017). However, as visualized by UMAP, the expanded NK cells in daclizumab beta-treated individuals cluster in the same region as CD56^bright^ NK cells from placebo-treated individuals, not with CD56^dim^ NK cells (Figure 1C). Instead, the CD56^bright^ NK cells appear to have acquired key characteristics that could enhance their cytolytic activity with daclizumab beta treatment, including enhanced expression of Perforin and CD16. Further, the daclizumab beta-induced CD56^bright^ NK cells express other markers of maturity, including KIR2DL3, Siglec-7, CD38, and CD8. Together, these data are consistent with the idea that the CD56^bright^ NK cells induced by daclizumab beta treatment are activated and have acquired sufficient maturity to be cytolytic.

Prior studies have indicated that NK cells may play a role in MS pathogenesis in the absence of drug treatment. Two groups have reported that CD56^bright^ NK cells are present in the CNS in MS lesions (B. Bielekova et al. 2011; Gross, Schulte-Mecklenbeck, Rünzi, et al. 2016), suggesting they may play a role in eliminating activated CD4+ T cells in brain lesions. Martinez-Rodriguez et al. reported that CMV infection, which drives an expansion of mature, ‘adaptive’ NKG2C-expressing NK cells, is associated with a lower risk of disease progression in MS (Martínez-Rodríguez et al. 2016). The CD56^bright^ NK cells observed after daclizumab beta do not resemble the mature NKG2C-expressing NK cells seen after CMV infection, but nonetheless could contribute to protection within the CNS.

Gross et al. report that CD56^bright^ NK cells are present in CNS lesions in MS, but that NK cells from MS patients are deficient in killing autologous, activated CD4+ T cells (Gross, Schulte-Mecklenbeck, Rünzi, et al. 2016; Gross, Schulte-Mecklenbeck, Wiendl, et al. 2016). This defect is attributed to poor expression of DNAM-1 on NK cells and its ligand, CD155 on CD4+ T cells in the setting of MS (Gross, Schulte-Mecklenbeck, Rünzi, et al. 2016), and is consistent with studies revealing that DNAM-1 polymorphisms may play a role in susceptibility to MS (Hafler et al. 2009). Independently, CD56^bright^ NK cells have been reported to mediate killing through NKG2D, TRAIL, and LFA-1 expression (Nielsen et al. 2012). In our study of peripheral blood NK cells, DNAM-1 expression was predictive of baseline samples, indicating that DNAM-1 expression is not increased by daclizumab beta treatment. However, it is important to note that our study characterized peripheral blood NK cells, and the prior studies showing DNAM-1-mediated killing were all focused on NK cells in the CNS.

Several studies suggest that NK cells may be deficient in the setting of MS (Benczur et al. 1980; Kastrukoff et al. 1988; French and Yokoyama 2004; Oger et al. 1988; Hauser et al. 1981; Merrill et al. 1982; Neighbour, Grayzel, and Miller 1982), providing hope that enhancing their frequency and/or function could improve outcomes. However, some studies have not demonstrated a defect in NK cell numbers in MS subjects (Gross, Schulte-Mecklenbeck, Rünzi, et al. 2016; Laroni et al. 2016). Another study suggests that ‘regulatory’ NK cells may be more important in disease pathogenesis (Aranami, Miyake, and Yamamura 2006). A limitation of our study was that our control group of healthy controls was collected independently of the SELECT and DECIDE trials. For comparison of CD56^bright^ frequencies, we used local healthy blood bank control subjects, but the potential for batch effects based on collection of blood samples at different times and with different methods (e.g. CPT tubes vs. heparin tubes), precludes our ability to compare NK cells between RMS subjects and healthy controls without concern for batch effects.

There are several limitations to our study. The first is that the sample size is quite modest, leading to reduced power to find differences in this high-dimensional analysis, particularly in the cross-sectional comparison between daclizumab beta-treated and placebo-treated individuals. Second, as discussed above, we did not have a healthy control group to which we could directly compare NK cells in RMS patients without concern for batch effects. Third, we did not perform functional assessments, so cannot confirm that the enhanced expression of killing and activation markers on CD56^bright^ NK cells in fact drives better killing of autologous, activated CD4+ T cells.

Overall, these data demonstrate that significant changes occur in NK cells in response to the increased IL-2 availability induced by daclizumab beta treatment. These data extend prior findings indicating that daclizumab beta treatment increases the frequency of the CD56^bright^ population, highlighting the unique phenotypic features of these expanded NK cells. The high expression of activation markers, activating receptors, and Perforin could enhance the ability of these CD56^bright^ NK cells to control RMS through cytolytic activity or other immunoregulatory functions. The deep profiling of NK cells performed here, including in the placebo group, can also serve as a reference for future studies of NK cell phenotype in the setting of RMS. While daclizumab beta was removed from the market due to serious adverse events, it is notable that several other medications used to treat RMS, including natalizumab, fingolimod, glatiramer acetate, or beta interferon, are also associated with expanded CD56^bright^ NK cells in the setting of clinical response (Caruana et al. 2017). Thus, in order to improve treatment for RMS, it will be critical in future studies to determine whether specific features of NK cells are associated with clinical response or serious adverse events.

## Acknowledgements

We thank Holden Maecker and Michael Leipold at the Human Immune Monitoring Core (HIMC) at Stanford University for use of their Helios machines and for their technical expertise. We also thank Wanda Castro-Borrero and Timothy Zheng at Biogen for their help compiling patient data and selecting samples for our study.

## Author contributions

CAB and JDF conceptualized this study. TR and CB designed experiments. TR and EV conducted experiments. TR, LS, and NZ analyzed the data with statistical analysis input from CS, AF, and SH. LS, TR, and CB wrote the manuscript; all authors contributed revisions.

## Supplementary Material

*Table S1. NK antibody panel for mass cytometry*

*Figure S1. Study design and characteristics of NK cells*

**(A)** Schematic of the study design. **(B)** Serial negative gating strategy used to identify total NK cells and CD56^bright^ and CD56^dim^ populations. Flow plots are from one representative individual. NK cells were isolated by magnetic bead purification prior to mass cytometry. **(C)** UMAP visualizations of total NK cells from the placebo treatment group at baseline. Each plot is colored by marker channel, with a color scale specific to each marker. CD56, CD16, NKG2A, CD57, NKp30, and Perforin were chosen to indicate where major NK subsets are localized in the UMAP visualizations of total NK cells.

*Figure S2. Example staining of all markers*

Each of the 31 NK markers in the antibody panel are shown against CD56 on the y-axis. Total NK cells were gated. Representative plots from one individual were chosen.

*Figure S3. Few differences between the placebo and daclizumab beta treatment groups at baseline*

A generalized linear model with bootstrap resampling was used to identify NK markers on total NK cells predictive of daclizumab beta- and placebo-treated individuals at baseline. Total NK cells were used with subsampling to 1000 cells per individual.

