## Supplementary Figures for "Characterization of the impact of daclizumab beta on circulating natural killer cells by mass cytometry"

Figure S1

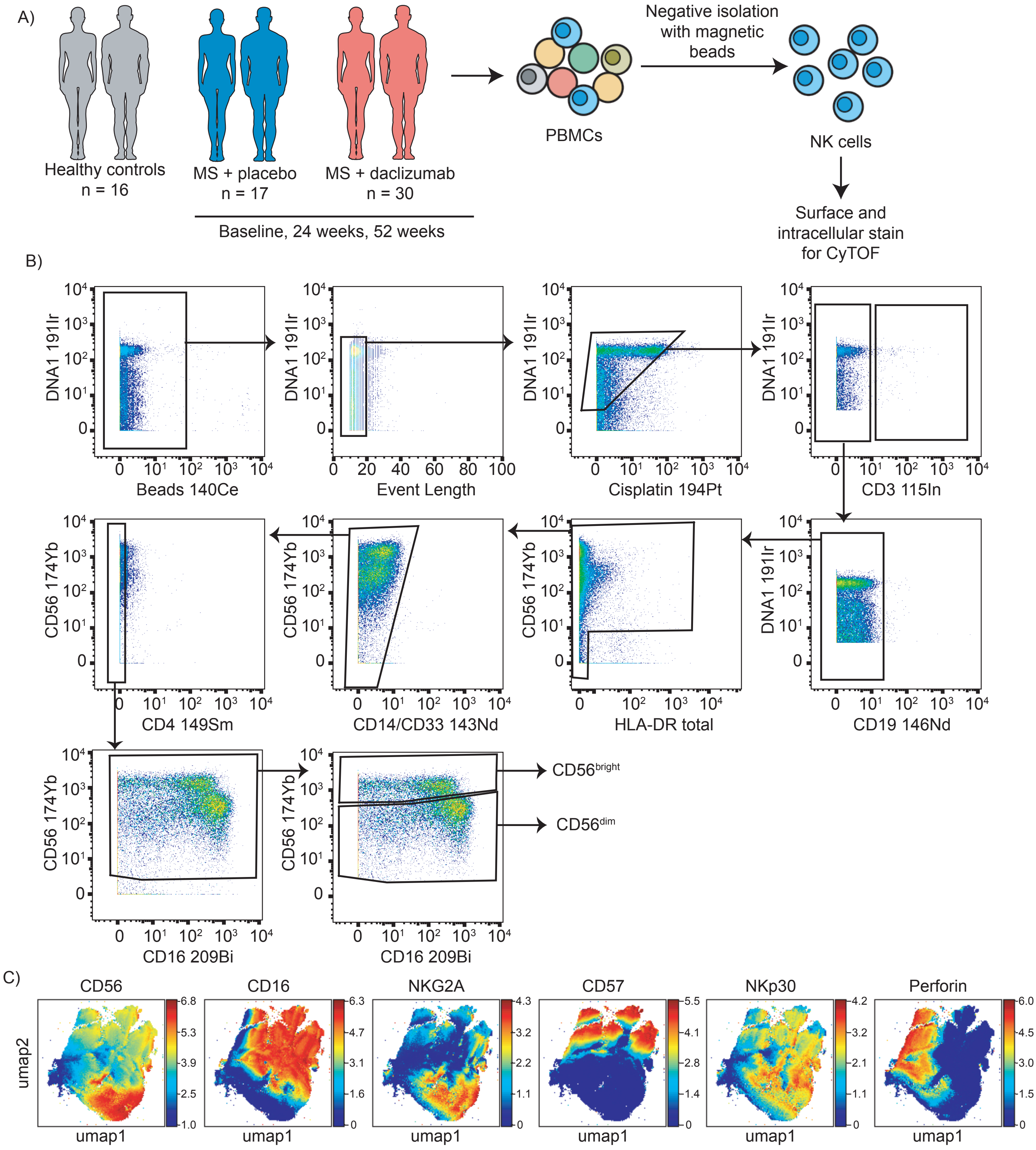


*Figure S1. Study design and characteristics of NK cells*

(A) Schematic of the study design. (B) Serial negative gating strategy used to identify total NK cells and CD56bright and CD56dim populations. Flow plots are from one representative individual. (C) UMAP visualizations of total NK cells from the placebo treatment group at baseline. Each plot is colored by marker channel, with a color scale specific to each marker. CD56, CD16, NKG2A, CD57, NKp30, and Perforin were chosen to indicate where major NK subsets are localized in the UMAP visualizations of total NK cells.

Figure S2

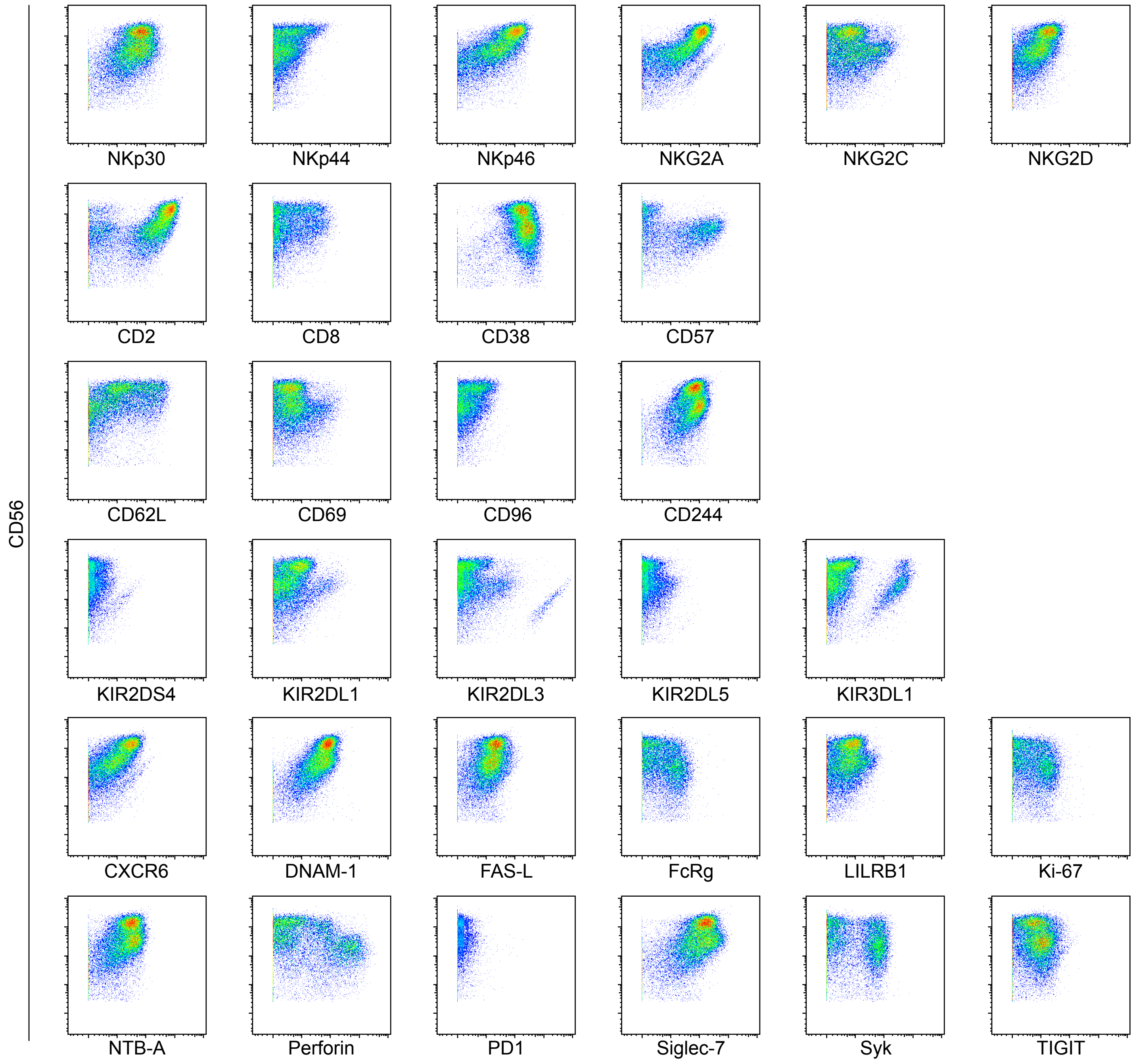


*Figure S2. Example staining of all markers*

Each of the 31 NK markers in the antibody panel are shown against CD56 on the y-axis. Total NK cells were gated. Representative plots from one individual were chosen.

Figure S3

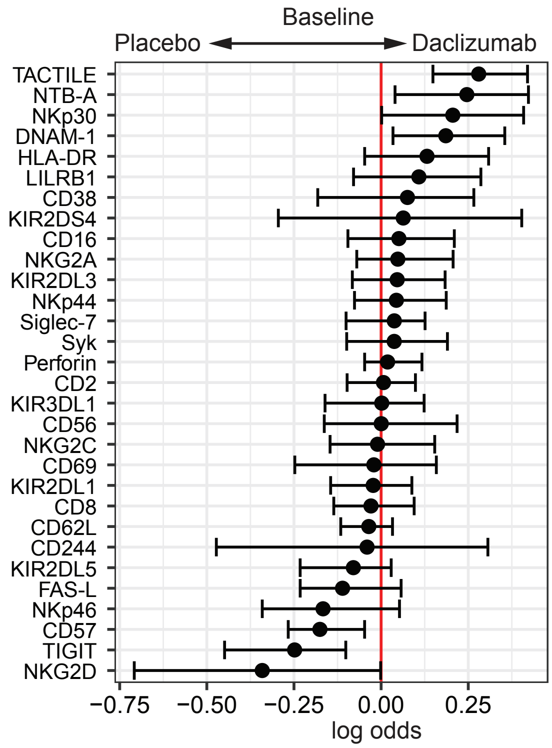


*Figure S3. Few differences between the placebo and daclizumab beta treatment groups at baseline*

A generalized linear model with bootstrap resampling was used to identify NK markers on total NK cells predictive of daclizumab beta- and placebo-treated individuals at baseline. Total NK cells were used with subsampling to 1000 cells per individual.
