## Supplementary Table 1 for "Characterization of the impact of daclizumab beta on circulating natural killer cells by mass cytometry"

**Table S1: NK antibody panel for mass spectrometry**

| **Isotope** | **NK Panel** | **Source** | **Clone** |
| --- | --- | --- | --- |
| 89Y | CD57 | Biolegend | HCD57 |
| Qdot | HLA-DR | Life Technologies | Tu36 |
| 115In | CD3 | Biolegend | UCHT1 |
| 141Pr | CD38 | Biolegend | HIT2 |
| 142Nd | CD69 | Biolegend | FN50 |
| 143Nd | CD33 | Biolegend | WM53 |
| 143Nd | CD14 | Biolegend | M5E5 |
| 144Nd | CD2 | Biolegend | RPA-2.10 |
| 145Nd | LILRB1 | Biolegend | GHI/75 |
| 146Nd | CD19 | Biolegend | HIB19 |
| 147Sm | CD8 | Biolegend | SK1 |
| 148Nd | FcRɣ | Millipore | Polyclonal |
| 149Sm | CD4 | Biolegend | SK3 |
| 150Nd | Syk | Biolegend | 4D10.2 |
| 151Eu | CD62L | Biolegend | DREG-56 |
| 152Sm | Ki-67 | Biolegend | Ki-67 |
| 153Eu | KIR2DS4 | R&D Systems | 179315 |
| 154Sm | KIR2DS2 | Abcam | Polyclonal |
| 155Gd | NKp46 | Biolegend | 9E2 |
| 156Gd | NKG2D | Biolegend | 1D11 |
| 157Gd | TIGIT | R&D systems | 741182 |
| 158Gd | 2B4 | Biolegend | C1.7 |
| 159Tb | DNAM-1 | BD Biosciences | DX11 |
| 160Gd | FAS-L | Biolegend | NOK-1 |
| 161Dy | NKp30 | Biolegend | P30-15 |
| 162Dy | Siglec-7 | Biolegend | S7.7 |
| 163Dy | NKG2C | R&D Systems | 134522 |
| 164Dy | NKp44 | Biolegend | P44-8 |
| 165Ho | CD96 | Biolegend | NK92.39 |
| 166Er | KIR2DL1 | R&D Systems | 143211 |
| 167Er | CD94 | Biolegend | DX22 |
| 168Er | CXCR6 | Biolegend | K041E5 |
| 169Tm | PD1 | Biolegend | EH12.2H7 |
| 170Er | KIR2DL5 | Miltenyi | UP-R1 |
| 171Yb | NKG2A | R&D Systems | 131411 |
| 172Tb | NTB-A | Biolegend | NT-7 |
| 173Yb | KIR3DL1 | BD Biosciences | DX-9 |
| 174Yb | CD56 | BD Biosciences | NCAM16.2 |
| 175Lu | KIR2DL3 | R&D Systems | 180701 |
| 176Yb | Perforin | Abcam | B-D48 |
| 209Bi | CD16 | Fluidigm | 3G8 |
